# Ecological and evolutionary drivers of thermal performance curves across the tree of life

**DOI:** 10.64898/2026.09.04.749418

**Authors:** Arnaud Sentis, Jérôme Eschenbrenner, Samraat Pawar, Anthony I. Dell

## Abstract

Biological rates typically follow a unimodal thermal performance curve (TPC), yet the ecological and evolutionary drivers shaping TPC variation remain poorly understood. We compiled an extensive thermal response database covering 896 TPCs for 416 species and 60 traits, spanning microbes to multicellular organisms. Our analyses reveal three key insights. First, environmental seasonality strongly structures TPCs: species in stable environments exhibit narrower thermal ranges and greater asymmetry, with environmental temperature outperforming latitude as a predictor. Second, trait type and organismal complexity modulate thermal sensitivity: emergent traits have narrower ranges than physiological traits, and complex organisms display reduced heat tolerance. Third, prokaryotes and eukaryotes differ fundamentally: while prokaryotes shift thermal limits symmetrically, eukaryotes show constrained upper limits relative to cold tolerance. These findings highlight how environmental stability, trait organization, and evolutionary history shape species’ thermal niches, providing crucial insights into vulnerability and adaptation under climate change.

## Introduction

Climate change profoundly impacts ectothermic organisms, affecting processes from intra- cellular mechanisms to interactions among individuals (1–4). The biological effects of rising temperatures on individuals and populations largely depend on how temperature influences biological rates from cellular to birth and death processes. Therefore, deepening our understanding of the thermal sensitivity of biological rates is crucial for accurately assessing and predicting the impacts of climate change on biodiversity.

The temperature dependence of biological rates follow a unimodal, asymmetric curve, referred to as Thermal Performance Curve (TPC, 4, 5, Fig. 1a). The TPC enables the examination of key physiological, behavioral, or ecological functions of a species across a broad temperature range. It can be measured for individual phenotypic traits, acknowledging that different aspects of species functioning respond uniquely to temperature changes (3). While the influence of temperature on phenotypic traits have stimulated decades of research (6–8), the ecological implication of TPCs in the context of climate change have only recently been thoroughly explored (4, 5, 9). Unlike vulnerability assessments focused solely on a species’ proximity to its thermal limits (10–13), the TPC approach incorporates the full temperature range, including non-lethal temperatures. This focus on the full temperature range is important because it allows the sensitivity of organisms to gradual warming within their “operational” temperatures to be quantified, thus providing a more balanced analysis of both positive and negative effects of climate warming. Understanding the factors influencing the key parameters of the TPC– minimal, optimal and maximal temperatures – is thus key to determining the thermal impacts on individuals and population and has been a central research focus in recent decades (2–4), from conservation (13, 14) to theoretical modeling (15).

**Figure 1:**
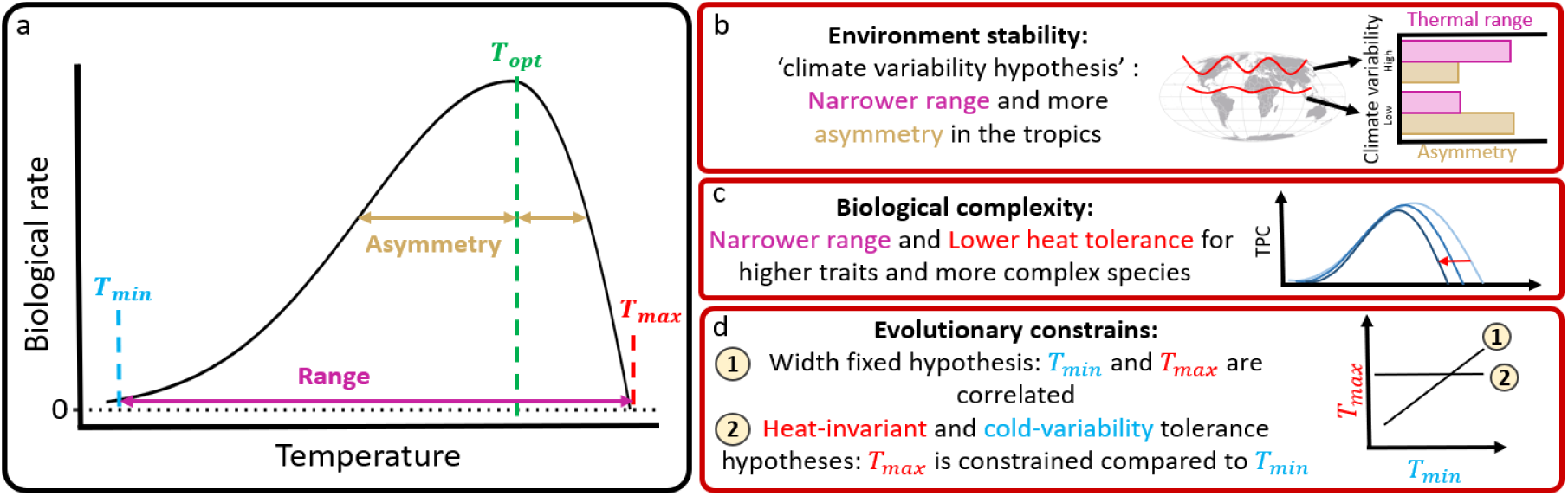
(a) A Thermal Performance Curve (TPC) describes how a biological trait responds to temperature and typically follows a unimodal and asymmetric bell curve. Key parameters are *T_opt_* (temperature at which biological rate peaks), *T_min_* and *T_max_* (the lower and upper thermal limits where rate reaches a minimum, respectively), thermal range (*T_max_*– *T_min_*), and asymmetry (difference in range between the rising and falling phases of the curve: (*T_max_* − *T_opt_*)/(*T_opt_* − *T_min_*) which also reflects the relative sensitivities (i.e. arrhenius activation energy) of these two phases). (b) Early theories such as the ‘climate variability hypothesis’ (Stevens 1989) examined how environmental fluctuations influence TPC parameters. (c) Trait and species complexity are likely to shape TPCs, yet their effects remain underexplored. (d) Finally, evolutionary constraints are expected to influence TPC shape, limiting the extent of thermal adaptation.

Environmental stability has been proposed as a key factor influencing TPC position and shape (16, 17). Climatic variability generally increases with latitude, giving rise to a widely discussed pattern: species at higher latitudes tend to exhibit a broader thermal tolerance range (5, 10, 13, 18–20), a pattern named the Janzen’s rule sensu Gaston et al. 2009. Following this idea, the “climate variability hypothesis” (21, Fig. 1b) proposes that a species’ thermal range is primarily influenced by the temperature fluctuations in its environment. Thermal ranges are expected to be narrower in environments with less temperature variability, as maintaining the physiological mechanisms necessary for a broad thermal range is energetically costly. Consequently, species in more stable, aseasonal environments should exhibit narrower thermal ranges (5, 10, 19). This hypothesis implies two macroecological patterns. First, because terrestrial environments are, on average, more thermally variable than aquatic environments (11, 13), thermal ranges should be wider in terrestrial than in aquatic species (13). Second, tropical species should have a narrower thermal range than temperate ones because temperature seasonality is weaker toward the equator (5, 10, 18, 19, 22). Third, tropical species should have more asymmetric TPC than those of temperate species (2, 19), because temperatures seldom exceed the optimal temperature for tropical species, thus their optimal temperature tends to be closer to their maximum heat tolerance (2, 18–20). Altogether this suggests that the thermal range of species and the degree of asymmetry in the TPC should reflect adaptation to environmental stability.

Another leading hypothesis is that the complexity of the organism or trait also drives the shape and position of TPCs. For organisms, complexity increases from simpler prokaryotes, which have less specialized cellular structures, to more advanced eukaryotes, which have organized organelles, and metazoans, which are multicellular animals with specialized organs and systems (23). Similarly, for traits, complexity refers to those traits that involve multiple interconnected physiological, biochemical, or morphological processes that require coordination among various components to function efficiently. Pörtner (24) proposes that more complex organisms (e.g., prokaryotes vs. eukaryotes vs. metazoans) should exhibit lower heat tolerance (Fig. 1c), as their intricate physiology (e.g., biochemistry) is more susceptible to disruption at higher temperatures. Building on the idea that thermal range correlates positively with organismal complexity, it is also expected that thermal ranges will be narrower for more emergent and complex integrated functional traits (3, 9, 24, Fig. 1c), as complex traits are inherently dependent on and embedded within simpler traits. For instance, Sentis et al (25) reported that respiration rate has a larger thermal range than consumption rate. If our prediction is correct, it implies that climate change might impact interaction traits differently than physiological traits(26).

Finally, evolutionary constraints likely shape TPCs, as tolerance for a wider range of temperatures is metabolically costly (27). Previous studies have argued for a tradeoff between adaptation to heat and cold, which would result in a fixed thermal range (25, Fig. 1d). While this relationship has not been widely reported (i.e. 26), it generally applies to certain taxonomic groups, such as ladybeetles (29). In contrast, several empirical studies spanning from terrestrial ectotherms (10) to plants (30) have found that heat tolerance tends to be relatively fixed across taxa, while cold tolerance is more flexible (5, 10, 12, 19, 22, 28, 29, Fig. 1d). In many cases, heat tolerance has shown limited capacity for adaptation (33). This observation led to the heat-invariant and cold-variability tolerance hypothesis, also known as the concrete ceiling, plastic floor hypothesis (16), first proposed by Brett (7).

While previous studies on TPCs have advanced our understanding of the factors influencing the shape and position of TPCs, they most often focus on a single driver (e.g. most often, latitude) or on specific taxonomic group (mostly terrestrial multicellular species). As a result, we lack a comprehensive view on the ecological and evolutionary mechanisms explaining variations in the shape and position of TPCs across latitude, habitat and taxonomic groups. To fill this gap, we compiled an extensive database of biological rates across temperatures from which we estimated 896 TPCs of ectotherms from a diverse range of species and traits. We then tested the main hypotheses detailed above that we grouped in three categories: environment stability (Fig. 1b), biological complexity (Fig. 1c) and evolutionary constraints (Fig. 1d). The taxonomic and trait diversity of our TPCs (Fig. 2a) allows us to explore the rarely tested co-effect brought by both traits and species complexity (3, 9, 24, 25). Overall, our aim is to disentangle ecological variations among groups and to reconcile opposing hypotheses, thus contributing to a comprehensive understanding of the dynamics associated with temperature-induced ecological changes. This knowledge is essential for accurately assessing species vulnerability and adaptation to changing temperatures.

**Figure 2:**
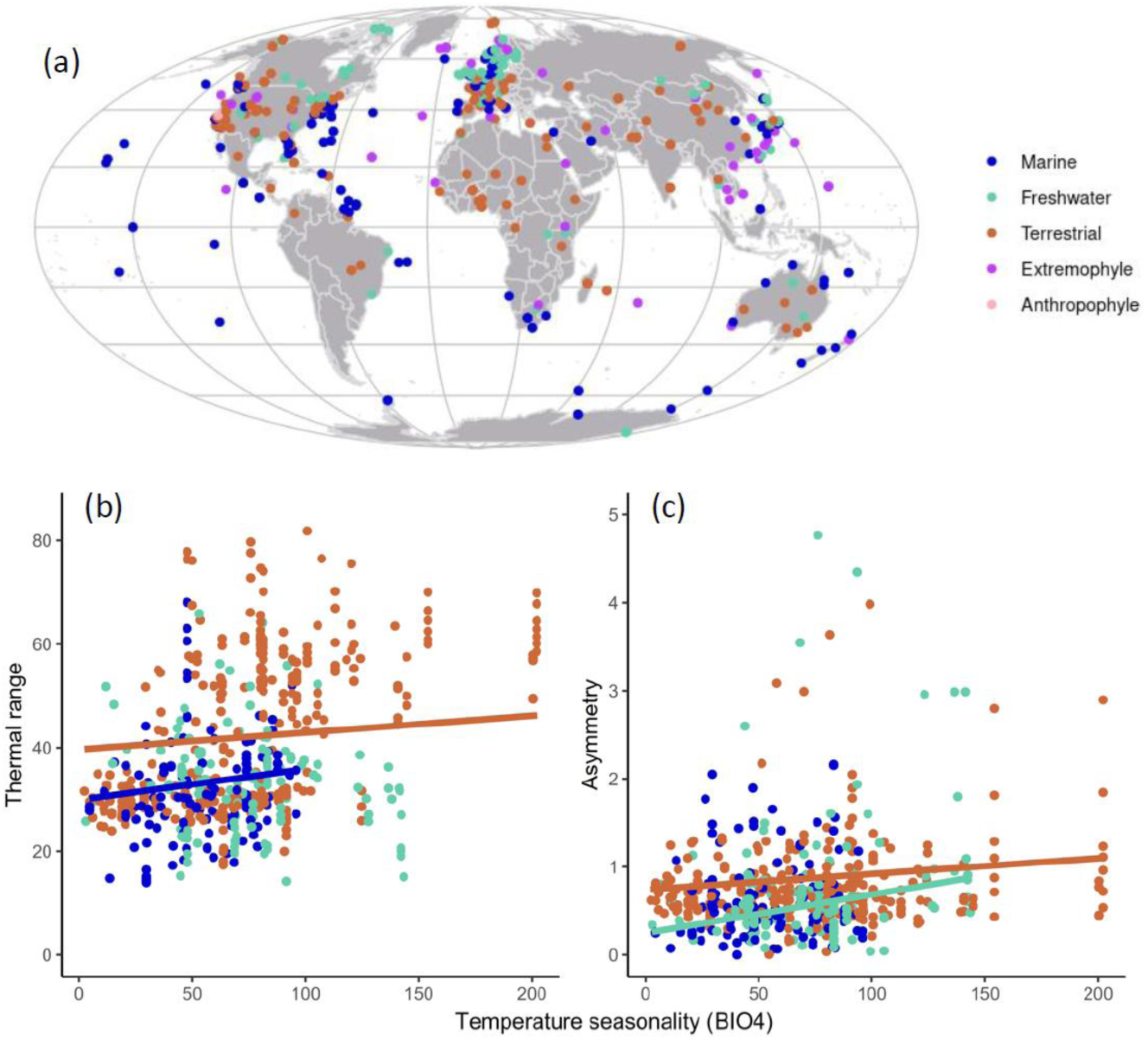
(a) map of the TPC data by habitat type. Relationship between temperature seasonality (WorldClim BIO4) and TPC thermal range (b) or asymmetry (c) for each habitat. Trend lines represent a significant relationship.

## Materials and Methods

### Data description

In a previous study (4), we gathered experimental data of biological rates at different temperatures from existing databases (such as Biotraits, 37) and datasets of other studies (such as 33), from prokaryotes (41) to multicellular ectotherm species (42), on numerous traits such as population growth, metabolism or inter-individual interactions. Our objective was to determine which model best fits the TPCs. In the present study we used these data to test the ecological and evolutionary factors influencing the shape and position of TPCs. The data are widely spread throughout the globe, taxa, and ecosystems (Fig. 2a). We selected data that included the following mandatory information: species name, measured trait name, trait values, and the associated temperature at which it was measured. Supplementary data, such as latitude or habitat, were retained when available. However, additional relevant data, such as species’ body size, individual age, or functional group (e.g., primary producer, predator), were insufficiently provided across TPCs for satisfactory analyses and are therefore included in SI for further reference.

We selected trait responses that were measured across a thermal range large enough to capture the rise and fall of biological traits with temperature and thus ensure correct estimates. For this purpose, we used four criteria: (i) having nonzero measurements for at least five distinct temperatures to fit our TPC models (some of them included 5 free parameters, see below); (ii) being hump-shaped: using the small-sample AIC (43), we selected TPCs better described by a quadratic polynomial model than by a linear one; (iii) being concave and having at least one temperature below and one temperature above the optimal temperature estimated from the quadratic linear model (we obtained the same selection of datasets if we choose datasets with two values above and two values below the optimum). This criteria allowed us to select TPCs describing the full temperature range of response traits and for which we had enough data to estimate TPCs.

### Fitting TPC models

A wide diversity of TPC models have been developed and although they perform differently, not a single model consistently performs better than others as we recently reported (4). In this study, we fitted 16 non-linear models to our temperature data, using a modified version of the R package rTPC (41, SI). Based on a preliminary screening, we found that Gaussian models best fitted our data. We thus fully fitted five models, which were the modified Gaussian model (45), the Pawar model (46), the Sharpe-Schoolfield high model (8) and the Deutch model (5). The Deutch model was the best fitting model, based on small-sample AIC and R² (SI Fig. S2). It is a bipartite model with four parameters, adapted from Deutsch (5) as:

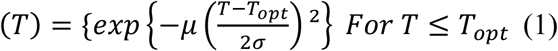

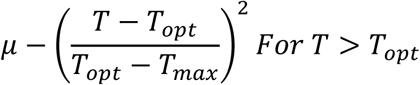

With *T_opt_* The optimal temperature at which the curve reaches its maximum, *σ* the variance of the Gaussian first half of the equation, *T_max_* the critical thermal temperature at which the function reaches 0 above the optimal temperature, and *μ* the maximum curve height at the optimal temperature. Using this model, we estimated several properties of the thermal window (Fig. 1a). The optimal temperature (*T_opt_*) and the thermal maximum (*T_max_*) are parameters of the Deutch model that were obtained directly from the model fit. Following Deutsch et al. 2018, we estimated *T_min_* as *T_opt_* − 4*σ*, where performance reaches a low value (as in Deutsch et al. 2008). We then defined the asymmetry of a performance curve as the ratio of its width on the warm and cold sides of *T_opt_*, so that asymmetry is equal to (*T_max_* − *T_opt_*)/(*T_opt_* − *T_min_*). The thermal range is defined as the difference between *T_min_* and *T_max_*.

We removed TPCs for which parameter estimates were very uncertain with standard error of *T_max_*, *T_min_* or *T_opt_* above 6°C (the standard error of *T_min_* was calculated using the law of propagation of uncertainty), or an R² under 0.4 or biologically not realistic with *T_max_* above 125°C or *T_min_* under -40°C. We also removed TPC with overly negative performance values (if |*min*(*rate*)| ≥ 0.15 *max* (*rate*)), as we could not estimate the base activity rate of those TPCs. We obtained a total of 896 TPCs, for 416 different species, 60 different traits with 459 unique combinations of species and traits.

### Reconstruction of the phylogeny

We included phylogeny in our analyses to account for the correlation between closely related species. As we did not find existing phylogenetic trees including all the 416 species, we reconstructed phylogeny using online tools based on the NCBI data. Because of the high taxonomic diversity across the 416 species, we were unable to gather satisfactory data on branch length and opted for a categorical approach using the *myTAI* dataset (47), and the *Open Tree of Life* project (48) for synonyms. Because taxonomic groups’ evolution time frames differ strongly depending on the kingdom making lower taxonomic levels (family, order…) incomparable (49), we only included the phylum and the class as a random effect.

### Environmantal drivers and catebogies

We gathered a proxy of measures of environment stability from Worldclim (Fick and Hijmans 2017) using spatial coordinates of each study and separating the species by their habitat as either “terrestrial”, “marine” or “freshwater”. Temperature seasonality (standard deviation of temperature, the BIO4 variable of Worldclim, AIC = 4136) was more strongly correlated to thermal ranges (*T_max_*– *T_min_*) than latitude (AIC = 4841). The temperature annual range (maximum – minimum temperatures, BIO7, AIC = 4137) was slightly less correlated than seasonality (BIO4).

We separated traits into three levels: “physiological” (such as metabolism), “emergent” (such as population growth) and “inter-individual interaction” (such as predation). We separated species complexity into three categories: “prokaryote”, “protist” and “multicellular eukaryote”. For trait complexity, the emergent category was by far the group with the most TPCs as it included all the measures of population growth (the most commonly measured trait). In contrast, we obtained very few inter-individual interaction TPCs, and these were largely limited to metazoans. We thus removed inter-individual interaction TPCs from our analyses of complexity. Too few physiological TPC were measured on protists; we did not remove them from the analyses to keep the category for others, but we did not interpret them for the protists.

### Statistical analysis

To test our hypotheses on the impact of environment stability, we investigated the influence of two categories of variables and their interactions on the thermal range and the asymmetry of TPCs. The first category is the habitat of the species, classified as “marine”, “freshwater” and “terrestrial”. Two more habitats, “extremophile” (e.g. hot spring or deep-sea vent) and “anthropophile” (living within human environment where the temperature is variable and unknown in much cases, such as microbes or food contaminant) were not represented in this analysis on environment stability as the temperature data we gathered did not reflect their local environment. The other category of variables represents annual temperature variation, including the commonly used latitude and two bioclimatic variables from WorldClim (50): temperature seasonality (standard deviation, BIO4) and temperature annual range (maximum – minimum temperatures, BIO7). We compared the AIC of models (using log-likelihood maximization) with habitat as a fixed effect and each of these three variables (and their interaction with the habitat), selecting the most parsimonious model, as the temperature in each habitat behave differently, with biological impact on thermal range (22).

To test our hypotheses on the impact of biological complexity, we examined how two of its indicators influence the thermal range of TPCs and the heat tolerance (*T_max_*) of species: (1) species complexity, classified as “prokaryote” (bacteria and archaea), “protist” (unicellular eukaryote) and “multicellular eukaryote” and (2) trait complexity, classified as “physiological” (such as cellular processes) “emergent” (resulting from a synergy of different traits, such as population growth) and “inter-individual interaction” (for interaction between individuals, sometimes different species).

To test the evolutionary hypothesis of the fixed thermal window width, we compared the variability in thermal range between species grouped by habitat and species complexity (and their interactions). As simply comparing the variance between those groups would not account for phylogenetic correlations, we transformed the data using mixed effect models with the phylogeny as random effects and extracting the residuals. Using these residuals, we could compare the variance between the groups using bootstraps on the entry data of the model (20 000 iterations) and compare the overlaps in the 95% confidence intervals (48, using the boot R package). To test the heat-invariant and cold-variability tolerance hypothesis, we compared the variance between species grouped by habitat and species complexity (and their interactions), using the residuals of two mixed effect models with both *T_min_* and *T_max_* as response variable and phylogeny as random effects. Using these residuals, we could compare the difference in variance within each group between both model (with *T_min_* and *T_max_* as response variable), using bootstraps (20 000 iterations) and compare the overlaps in the 95% confidence intervals.

All statistical analyses were performed using the package nLME (52) in R software (version 3.6.3, R development Core Team, 50), to perform mixed effect models with the phylogeny as a hierarchical random factor on the intercept. Deviance analyses were carried out based on likelihood ratio tests to determine the significance of the fixed effects. If there was a significant interaction among fixed variables, we performed post-hoc analyses by dividing the dataset by the variable of interest and fitting again the mixed model to the data subsets to understand all facets of interactions. Assumptions of each model were verified by checking the residuals.

## Results

### 1) Environment stability

Narrower thermal ranges in more stable environments. We gathered a proxy of measures of environment stability from Worldclim (Fick and Hijmans 2017) using spatial coordinates of each study and separating the species by their habitat as either “terrestrial”, “marine” or “freshwater”. Temperature seasonality (standard deviation of temperature, the BIO4 variable of Worldclim, AIC = 4136) was more strongly correlated to thermal ranges (*T_max_*– *T_min_*) than latitude (AIC = 4841). The temperature annual range (maximum – minimum temperatures, BIO7, AIC = 4137) was slightly less correlated than seasonality (BIO4). Both seasonality (*F=5.31, p=0.021*) and habitat (*F=5.85, p=0.003*) had a significant impact on thermal range, as well as their interaction (*F=4.61, p=0.01*). Marine species had the narrowest thermal range, followed by freshwater and then terrestrial species (Fig. 2b). Moreover, both marine and terrestrial species’ thermal ranges increased with temperature seasonality, but seasonality had no significant influence on freshwater species.

Increased TPC asymmetry in more stable environments. Species’ traits with a *T_opt_* closer to their *T_max_* than to their *T_min_* were considered to have a more asymmetric TPC. Temperature seasonality was more strongly correlated to TPC asymmetry (AIC = 934) than was latitude (AIC = 1435) or temperature annual range (AIC = 936), with latitude failing to explain any significant change in asymmetry. We observed a significant interaction between habitat and seasonality (*F=4.65, p=0.01*), suggesting that differences in asymmetry among habitats depend on seasonality, with marine species failing to show changes in asymmetry with seasonality.

The TPCs of freshwater species were more asymmetric than those of terrestrial species at the intercept (i.e., in environments with lower temperature variability, such as the tropics), with their asymmetry increasing more rapidly as seasonality increased (Fig. 2c). While terrestrial and marine species did not differ significantly in terms of intercept or rate of change with seasonality, the larger dataset for terrestrial species revealed a significant decrease in asymmetry with increased seasonality, and a notable difference in asymmetry compared to freshwater species.

### 2) Biological complexity

We separated traits into three levels: “physiological” (such as metabolism), “emergent” (such as population growth) and “inter-individual interaction” (such as predation). We separated species complexity into three categories: “prokaryote”, “protist” and “multicellular eukaryote”. For trait complexity, the emergent category was by far the group with the most TPCs as it included all the measures of population growth (the most commonly measured trait). In contrast, we obtained very few inter-individual interaction TPCs, and these were largely limited to metazoans. We thus removed inter-individual interaction TPCs from our analyses of complexity. Too few physiological TPC were measured on protists; we did not remove them from the analyses to keep the category for others, but we did not interpret them for the protists.

#### Complexity impact on thermal range

No significant differences in thermal range were found between species complexity (F = 1.70, p = 0.199), although multicellular species exhibited the widest range. Overall, the thermal ranges of emergent traits were consistently narrower than those of physiological traits (F = 7.348, p = 0.007, Fig. 3a). The interaction term was only significant due to the small protist sample size mentioned earlier and lost its significance when these data were excluded.

**Figure 3:**
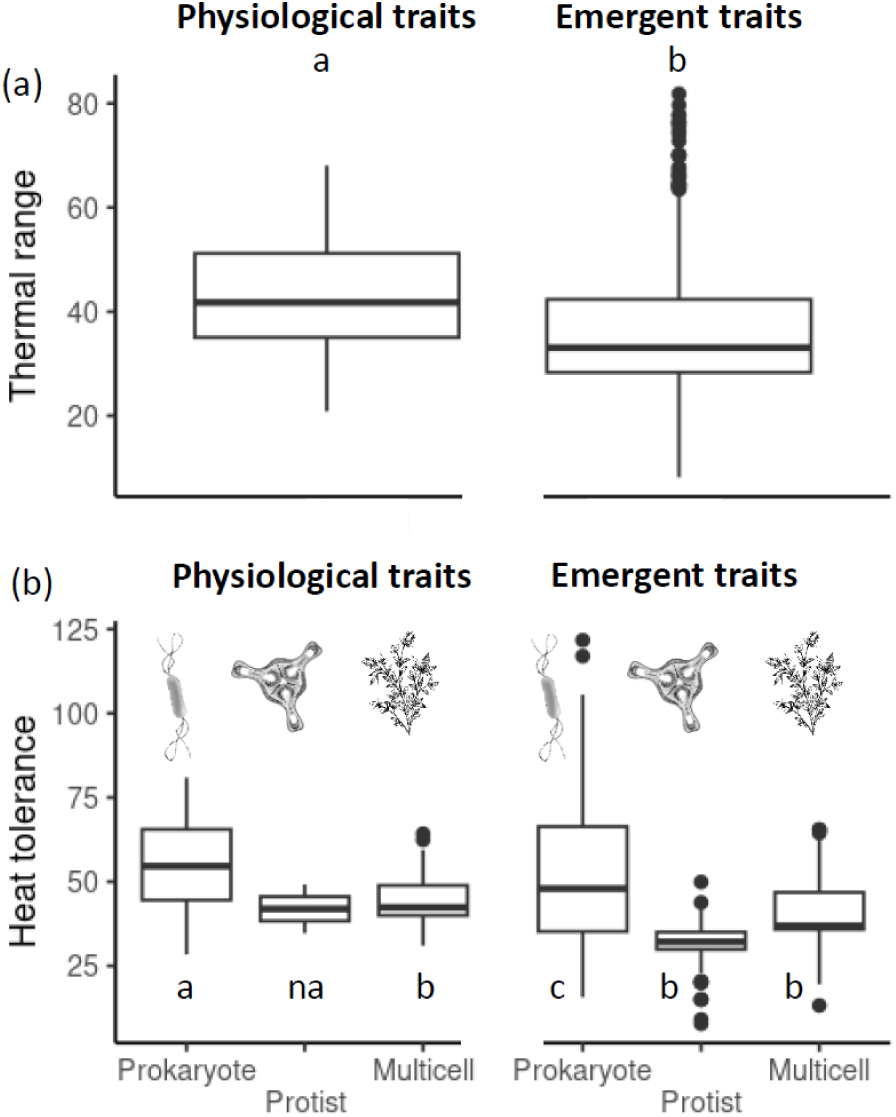
Thermal range (a) and heat tolerance (b) in function of trait and species complexity. Species complexity is not represented in (a), as it was not significant. Letters represent statistical significance.

#### Complexity impact on heat tolerance

Species complexity (*F=12.135, p=0.0018*), trait complexity (*F=14.729, p=00.0001*) and their interactions (*F=9.259, p=0.0024*) significantly influenced the heat tolerance of species (Fig. 3b). For both physiological and emergent traits, heat tolerance in eukaryotes was significantly lower than in prokaryotes, with no significant differences between protists and multicellular eukaryotes. For prokaryotes, heat tolerance was significantly higher for physiological traits compared to emergent traits, whereas, for multicellular species, no significant differences were observed between physiological and emergent traits.

### 3) Evolutionary constraints

#### Fixed-width hypothesis

A thermal range with a fixed width implies that *T_max_* and *T_min_* change together and are thus positively correlated. We tested this correlation with a linear model that included species complexity (other variables such as habitat and trait complexity only weakly influenced those correlations). While *T_min_* and *T_max_* were correlated (*F=592.5, p<0.0001,* Fig. 4a), the slope of the relationship depended on species complexity (*F=89.4, p<0.0001*); it was only strong for prokaryotes (1 *T_min_* for 0.79 *T_max_, 95% interval: 0.73- 0.85*), and weak for protists (1 *T_min_* for 0.25 *T_max_, 95% interval:0.12 - 0.39*) and multicellular eukaryotes (1 *T_min_* for 0.13 *T_max_*, *, 95% interval: 0.03 - 0.23*). In any case, all slope values were below one, suggesting no isometry between *T_min_* and *T_max_*.

**Figure 4:**
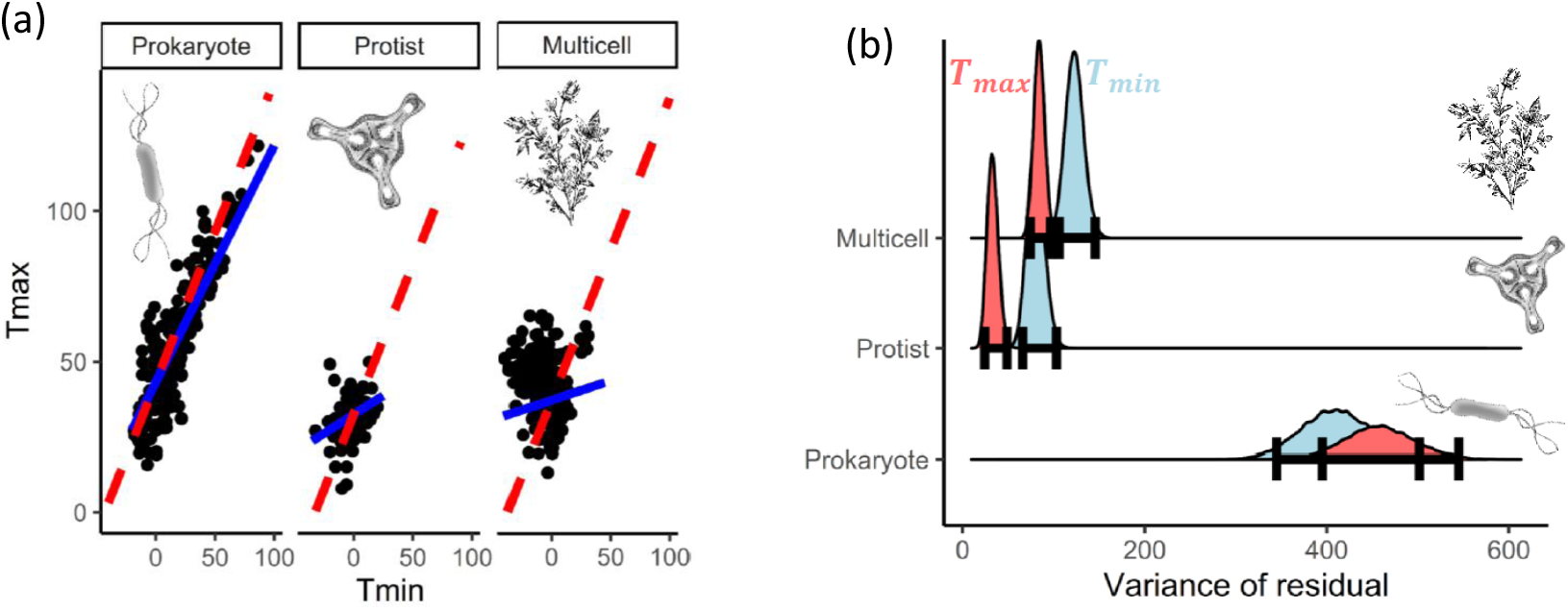
Test of both evolutionary constraint hypotheses on *T_min_* and *T_max_*, for each species complexity group. (a) Width fixed hypothesis based on the correlation between *T_min_* and *T_max_* . The red dotted lines represent the 1:1 line, the blue lines represent significant correlations. (b) Heat-invariant and cold-variability tolerance hypothesis. Variability of *T_min_* (blue) and *T_max_* (red) after correction by the phylogenetic random effects. The bold lines represent the 95% confidence interval.

#### Heat-invariant and cold-variability tolerance hypothesis

We compared variabilities of heat and cold tolerance for each species complexity level using the residuals of a mixed effect model to account for phylogeny, and used bootstraps on the variability to test for significance. *T_max_* was less variable than *T_min_* for multicellular species (*95% interval: T_min_ 106.0 - 145.8, T_max_ 74.1 - 96.5*) and protists (*95% interval: T_min_ 65.8 - 103.1, T_max_ 24.2 - 48.7*), whereas, for prokaryotes, variability was not significantly different for *T_min_* and *T_max_* (*95% interval: T_min_ 344.9 - 501.8, T_max_ 395.2 - 545.1*, Fig 4b). However, when separating by habitat, among the multicellular species, only the terrestrial ones had a less variable *T_max_*, and marine prokaryotes also differed from other prokaryotes by having a significantly less variable *T_max_* (supporting information (SI) Fig. S3). Adding or removing the inter-individual interaction trait, which was only measured in multicellular species, did not alter our results.

## Discussion

Previous studies on TPCs have improved our understanding of the factors shaping TPCs, but they have typically focused on a single hypothesis (e.g., latitude) or specific taxonomic groups (primarily terrestrial multicellular species). In this study, we present the first comprehensive analysis of the ecological and evolutionary mechanisms driving variations in the shape and position of TPCs across different latitudes, habitats, and taxonomic groups. Our results indicate that no single hypothesis is universally valid; instead, the primary mechanisms driving TPC variations depend on the species’ habitat as well as the complexity of the trait or species.

### Environment stability

Our results indicate that species inhabiting more stable thermal environments, such as aquatic habitats or regions with lower climatic variability, tend to have narrower thermal ranges. This pattern aligns with Janzen’s rule (16) and other studies (11, 13), suggesting that species evolve narrower thermal tolerances in stable climates because there is less selective pressure to withstand a wide range of temperatures. In contrast, species in more variable environments must adapt to broader temperature fluctuations, resulting in wider thermal ranges.

The exception found in freshwater species highlights the unique thermal dynamics of these habitats. Unlike marine or terrestrial systems, freshwater environments often exhibit localized temperature conditions influenced by factors like water flow, shading, and proximity to sources(34). This may indicate that thermal adaptation in freshwater species may be driven by fine-scale, local temperature patterns rather than broad climatic trends. This also suggests that freshwater species are more specialized to their microhabitats, making them potentially more vulnerable to local disturbances or changes in thermal regimes.

Overall, our findings emphasize the role of environmental stability in shaping thermal performance curves (TPCs). Our results suggest that species in stable climates may face greater risks from climate change, as their narrower thermal ranges limit their capacity to cope with rising temperatures. In addition, these species might experience stronger selection pressures under rapid climate warming, potentially leading to shifts in their thermal optima or increased extinction risks if they cannot adapt quickly enough (35).

Regarding asymmetry, we found that the *T_opt_* of terrestrial and freshwater species shifts closer to their *T_max_* as temperature seasonality decreases. Since hotter environments typically exhibit lower variability, our findings align with Buckley (2), who observed greater asymmetry in such warmer environments. Several mechanisms are likely to play an important role in explaining this pattern. First, global seasonal variation strives for species to be acclimated to a variety of temperatures. Second, although tropical species experience high mean temperatures, the daily temperature maximum in these regions tends to have lower variability compared to temperate climates. This reduces the evolutionary pressure for species to develop mechanisms for additional heat tolerance. As a result, the optimal temperature of tropical species is closer to their maximum heat tolerance (2, 18–20), making them more susceptible to rising temperatures driven by climate change (2, 5, 19, 20). Third, from an evolutionary perspective, heat tolerance shows little variability (Fig. 4, 5, 10, 12, 19, 22, 28, 29) and low evolutionary potential (33). The optimal temperature is more variable (2, 5) and is expected to shift more rapidly toward warmer temperatures as the environment warms, thereby increasing asymmetry. However, unlike freshwater and terrestrial habitats, we found no significant differences in asymmetry with increasing seasonality for marine species. While some argued that marine species were less vulnerable to climate change (35), others found the opposite (13, 14). The greater vulnerability in those studies could arise from other factors not captured in our data, such as the limited availability of thermal refugia (13).

Our analysis shows that temperature variability, rather than temperature extremes or latitude, better explains differences in species’ thermal ranges. While latitude has traditionally been used as a proxy for thermal tolerance in many studies (2, 5, 10, 11, 13, 16, 22, 36, 37), it does not capture the finer-scale climate dynamics that influence thermal adaptation. Similarly, although the Climate Extremes Hypothesis (22, 38) posits that extreme temperatures are key in shaping thermal tolerances, our results indicate that overall temperature variability (BIO4) is a stronger predictor. The limited explanatory power of temperature extremes (BIO7) might reflect the fact that species often face a range of non-lethal temperatures that shape their performance curves more effectively than rare extreme events. These results emphasize the importance of considering temperature variability as a key factor in understanding species’ thermal adaptation and predicting their responses to climate change.

### Complexity

Our data provide good evidences supporting the hypothesis that the thermal range of emergent traits—such as population growth, locomotion, or feeding ability—is nested within more fundamental physiological traits, as previously proposed by Sentis (25), Bozinovic (32) and Ørsted (3). This pattern is likely due to the dependence of emergent traits on multiple physiological traits, which constrains the thermal range of emergent traits similar as in the Leibniz’s principle of minimum requirements which states that growth is controlled not by the total availability of resources, but by the scarcest one. In thermal biology, this means that an organism’s performance at higher levels of organization (e.g., population growth) is bounded by the physiological function that fails first under thermal stress. The ecological and evolutionary implications of this nested pattern are significant. Because emergent traits rely on multiple physiological traits, their thermal range tends to be narrower, limiting species’ ability to maintain key biological functions such as reproduction or foraging under thermal stress.

This could influence species’ resilience to climate change, as organisms may be unable to maintain optimal performance in warmer conditions if their physiological traits are already near their thermal limits. From an evolutionary perspective, the relationship between emergent and physiological traits suggests that evolutionary shifts in thermal tolerance might first be seen in more fundamental physiological processes, such as metabolism, before impacting emergent traits like population growth or behavior. Recent work on arthropods (26) supports this view, showing that the evolutionary capacity for thermal adaptation is constrained by life-history trade-offs and energetic limitations. Indeed, the peak performance temperatures of different physiological traits often differ and do not shift uniformly, likely due to conflicting physiological demands. Altogether, these findings highlight that both the performance and evolutionary response of emergent traits are tightly constrained by physiological limitations and trade-offs.

We also report the first evidence that more complex organisms have a lower heat tolerance. As suggested by Pörtner (24), this may be explained by more complex and interdependent metabolic processes in complex eukaryotes compared to simpler prokaryotes species. Eukaryotes did not exhibit a significant change in heat tolerance across trait complexity, which aligns with our subsequent analysis showing that their heat tolerance is generally constrained (but see 29). In contrast, for prokaryotes, heat tolerance decreased with higher trait complexity, which fits with the general pattern of emergent traits having more constrained thermal limit than physiological traits. The finding that more organized, complex organisms tend to have lower heat tolerance has important ecological implications. The constrained heat tolerance in eukaryotes aligns with the idea that their complex metabolic systems require more energy and are more susceptible to disruption under thermal stress, which may limit their ability to tolerate higher temperatures. Conversely, the observed decrease in heat tolerance with increasing trait complexity in prokaryotes supports the concept that emergent traits, which are often dependent on fundamental physiological traits, have more restricted thermal limits. This suggests that the vulnerability of species to climate change may be tightly linked to their physiological complexity, with more complex organisms potentially facing greater risks from thermal stress, as their intricate systems are more sensitive to temperature fluctuations.

### Evolutionary constraints

Previous studies have proposed two contrasting hypotheses regarding constraints on the shape of the TPCs. The fixed TPC width hypothesis suggests that *T_max_* and *T_min_* must change in tandem (27), whereas the heat-invariant and cold-variability tolerance hypothesis (7, 16) posits that changes in thermal range width occur because cold tolerance is more variable than heat tolerance. We found that the validity of these two hypotheses depends on phylogenetic groups. Only prokaryotes had a relatively fixed thermal range (in line with the fixed TPC width hypotheses), while terrestrial multicellular eukaryotes’ TPC thermal range was highly variable. In addition, only eukaryotes had a relatively fixed heat tolerance compared to their cold tolerance (in line with the heat-invariant cold-variability hypotheses). Overall, we found that both hypotheses apply to different species groups, making them mutually possible. This highlights the need to consider species-specific differences when working at a large scale, as broad comparisons offer new insights, but finer-scale studies are crucial for fully understanding each hypothesis.

However, classifying species based solely on their organization does not fully explain whether a species adheres to each hypothesis. For instance, marine species exhibited an inverse pattern to the heat-invariant and cold-variability tolerance hypothesis compared to other habitats (SI Fig. S3), and other studies, such as Dixon’s (29), have found results that do not align with our classification. It is therefore possible that another factor, such as a microhabitat or taxonomy interaction, governs the transition between these two hypotheses, with our classification serving as a proxy for this relationship. Moreover, previous research mostly focused on terrestrial multicellular species (5, 10, 12, 30, 31, 39) and our analyses for this group yield similar results. This focus on terrestrial multicellular species could explain the lacking literature comforting the width fixed hypothesis. On other species groups, these hypotheses have mostly not been tested, and our study provides a first test of the differences among terrestrial, marine and freshwater multicellular species. For instance, Sunday (22) showed that marine and freshwater multicellular species exhibited more variable heat tolerance than terrestrial species, a finding we also observed.

Testing the width-fixed hypothesis with our data presents limitations, as we can only assess correlations, which may arise from confounding factors. For instance, in environments with similar levels of seasonality, maximum and minimum temperatures are often correlated, making it difficult to isolate independent effects. However, the diversity within our dataset allows for a more nuanced examination of this hypothesis. Among prokaryotes, extremophiles — species adapted to extreme heat in environments such as hot springs or deep-sea vents — align particularly well with the width-fixed hypothesis. These species rely on high heat resistance for survival, yet despite the presence of nearby habitats with more moderate temperatures, their limited cold tolerance prevents them from colonizing those areas. This pattern suggests that the thermal range of their TPCs is evolutionarily constrained, reinforcing the idea that adaptation to extreme conditions imposes trade-offs on thermal tolerance.

## Conclusion

By merging diverse datasets, we were able to study the thermal performance curves of an unprecedented range of taxa and habitats. Our findings confirm the Climate Variability Hypothesis while revealing important differences across habitats. To refine our understanding, we recommend using temperature data instead of latitude because the latter is merely a proxy for temperature variation. Additionally, we demonstrate that greater biological complexity increases vulnerability to climate change by reducing both maximum temperature tolerance and thermal range. We also reconcile opposing hypotheses by showing that evolutionary constraints drive differences in thermal responses across taxonomic groups. Overall, our findings suggest that the impacts of climate change on traits and species are shaped by their environmental conditions and evolutionary history, providing crucial insights into species vulnerability and adaptation to rising temperatures.

## Acknowledgments

This work was supported by the ANR project EcoTeBo (ANR-19-CE02-0001-01) from the French National Research Agency (ANR) to AS.

## Notes

### Competing Interest Statement

The authors have declared no competing interest.

