## Supporting Information for "Ecological and evolutionary drivers of thermal performance curves across the tree of life"

### Modifications of the rTPC package

The rTPC package (Padfield and O'Sullivan 2021) provides 25 different functions harvested from the literature to fit experimental TPC data. We reused their source code and modified it when necessary so that it ran on more complex data such as with temperature duplicates. We removed the functions that did not have a hump shape, and the ones that failed to fit part of our data. The rTPC package can be found at: https://padpadpadpad.github.io/rTPC/index.html

### Choice of the best model for fitting TPC

Based on preliminary analyses where we fitted 16 models to our TPCs, we selected the best four fitting ones. They were the Gaussian model (Angilletta 2006), the Pawar model (Kontopoulos 2018), the Sharpe-Schoolfield – high model (Schoolfield et al. 1981) and the Deutch model (Deutsch et al. 2008). We selected the best model using $R^2$ and small-sample AIC (Fig. S2).

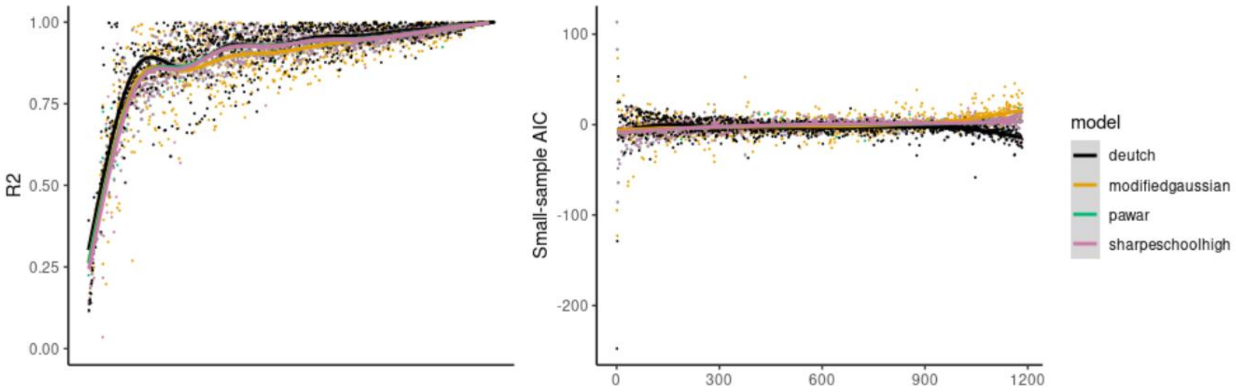


Figure S2: the fitting quality of our four models to each of our TPCs, quantified by (a) $R^2$ and (b) small-sample AIC. For better visualization, we ordered the x-axis by the mean of the y-axis between the models. To allow comparison between each model small-sample AIC, we centered the values of each model in-between each TPCs.

The mean $R^2$ of each model was 0.883 for the Gaussian model, 0.89 for the Pawar model, 0.887 for the Sharpe-Schoolfield – high model and 0.904 for the Deutch model, respectively. The mean difference of small-sample AIC compared to the other model was 1.693 for the Gaussian model, 0.132 for the Pawar model, 0.274 for the Sharpe-Schoolfield – high model and -2.098 for the Deutch model. The Deutch model was thus the best fitting model for our TPCs.

### Detailed graphics of the heat-invariant and cold-variability tolerance hypotheses by habitats

Due to differing sample sizes across habitats, terrestrial species were more frequently represented in our dataset. To address this, we conducted additional analyses that revealed considerable variability in the patterns observed between habitats for the heat-invariant and cold-variability tolerance hypotheses.

When we examined species complexity within each habitat, distinct patterns emerged (Fig. S3). Terrestrial multicellular species exhibited more variability in cold tolerance compared to heat tolerance. In contrast, freshwater and marine multicellular species did not show this pattern. Interestingly, freshwater species did not align with either hypothesis, suggesting unique tolerance dynamics in this environment. Marine species presented an inverse pattern relative to terrestrial species: while multicellular marine species did not support the hypotheses, marine protists and prokaryotes showed less variability in heat tolerance than in cold tolerance. These findings highlight the complexity and habitat-specific nature of thermal tolerance across different groups of organisms.

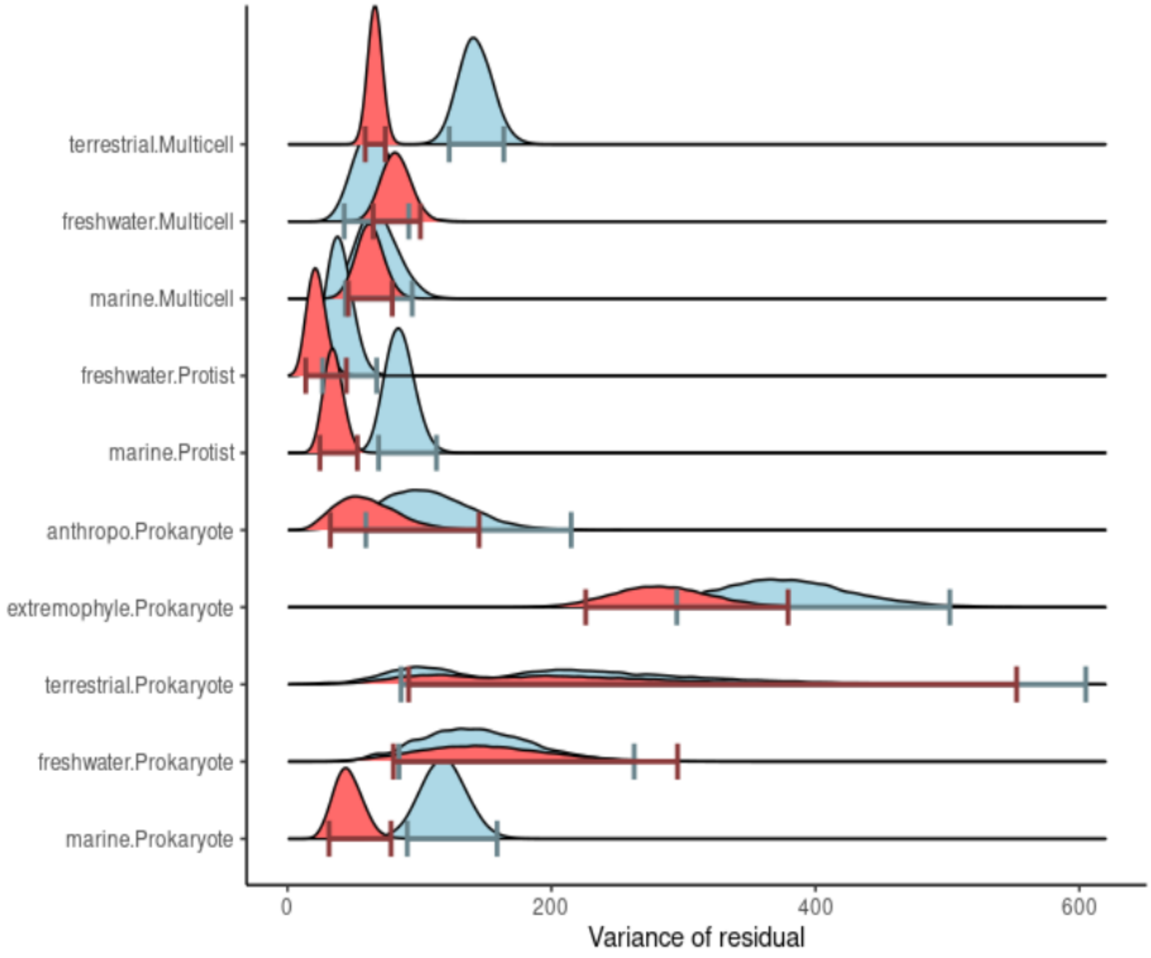


Figure S3: Comparison of the variance between the heat and cold tolerance in-between each group, to test the Heat-invariant and cold-variability tolerance hypothesis. Variance of the residual of mixed effect model with the phylogeny as random factor, separated by both habitat and species organization. Interval of confidence of the variance of the residual by 95%. Multicellular species and the habitat of anthropophile are shortened respectively to “Multicell” and “anthropo”.

### Size dependence of thermal window

Body size plays a significant role in shaping species' ecological strategies. Larger species are generally expected to tolerate a broader range of temperatures (e.g., Kaspari et al. 2015), as they possess more options for thermoregulation—including greater mobility and higher thermal inertia. Conversely, smaller species can exploit fine-scale microclimates, which may buffer them against extreme temperatures. These microclimates are also more commonly available in terrestrial ecosystems.

However, our dataset included only 175 species across the three major habitat types, which limited our ability to robustly test these hypotheses—particularly given the high phylogenetic diversity represented.

#### Ontogeny

In our dataset, we identified 160 samples taken from adult stages and 49 from juvenile stages, all belonging to multicellular species (plants, fungi, and metazoans). Unfortunately, very few species were represented by both life stages, limiting our ability to draw direct comparisons within species. As a result, we were unable to reach definitive conclusions. However, we did observe that thermal range variability appeared to be higher among juvenile samples compared to adults, suggesting potential differences in thermal tolerance across life stages that warrant further investigation.

#### Functional groups

Dixon et al. (2009) showed that, in an insect dataset, predators and parasitoids exhibited narrower thermal windows than other trophic groups. This suggests that species higher up the food chain may generally have more limited thermal ranges, potentially due to their dependence on the thermal tolerances of their prey. Unfortunately, we were unable to test this hypothesis in our study due to insufficient data across functional groups. Our dataset included only 141 carnivores, 12 omnivores, 22 herbivores, and 4 primary producers, which did not allow for meaningful comparisons.
